# Discovery of the spatial pattern of prickles on the stem of the rose and the mathematical model of the pattern

**DOI:** 10.1101/811281

**Authors:** Kazuaki Amikura, Hiroshi Ito

**Affiliations:** Department of Molecular Biophysics and Biochemistry, Yale University, New Haven, Connecticut, USA; Department of Computational Biology and Medical Sciences, Graduate School of Frontier Sciences, The University of Tokyo, Japan; Department of Art and Information Design, Faculty of Design, Kyushu University, Japan

**Keywords:** Plant development, Rose, Prickle, Mathematical model

## Abstract

Reproducible pattern is a key characteristic of organisms. Many developmental patterns are known that it is orchestrated by diffusion of the factors. Herein, we reported a novel patterning that seems to be controlled by diffusion factors. Although it looks like the prickles randomly emerge on the stem of rose, we deciphered patterns for the position of prickles with statistical data and proposed a mathematical model to explain the process via which the pattern emerged. By changing the model parameters, we reproduce another pattern on other plant species. This finding indicates that the patterns between many species are organized by similar systems. Moreover, although the pattern of organisms is often linked to its function, we consider the spatial pattern of prickles may have a function to play the role of prickles effectively. Further studies will clarify the role of prickles and reveal the entity of diffusive factors.

---

The pattern formation is fundamental characteristic of life. Many pattern formations are expected that it is orchestrated by diffusion of the factors. Gradient of diffused factors are considered key factor of development in early 20th century(1, 2). In 1952, Alan M. Turing provide conceptual formulated solution to explain the pattern formation based on the diffusion(3). For half a century from then, it has been shown that his model applies to many phenomena of the development of life(4, 5).

In the pattern formation of plant, the diffusion of auxin that is a plant hormone plays a central role(6). Especially, the development mechanism of the phyllotaxis pattern that is the positioning of leaves on the stem has well understood based on the diffuse of auxin. It is proposed that the phyllotaxis of *Arabidopsis* is dependent on polar auxin transport.(7, 8). The pattern of leaf venation is also explained by polar auxin transport in leaves(9). Otherwise, the position of stomatal and trichome are considered to be organized by diffusing factors such as protein and peptide(10). As these results show, the diffusion of the factors is fundamental phenomena for the pattern forming of plant.

This paper focuses on the pattern of prickles on the stem of rose. Although research about the flower reported from various viewpoints, but the prickle on the stem and the petiole is not well studied. Anatomical studied showed that the glandular trichome developed from an epidermal tissue seems to be grown into prickle in rose(11). Recently, the candidate gene that controls the density of prickle on the stem has been suggested(12), while the gene concerning the emergence of prickle is not revealed yet (13, 14). Therefore, the molecular mechanism of the development of prickle is not understood as well as other protrusion structures on the stem, such as thorn and spine. Moreover, the pattern of prickles on the stem of rose is not studied with statistical data and mathematical model in the long history of rose.

In this paper, we decipher the pattern of prickles on the stem of rose has the pattern and suggests the mathematical model based on the diffusion. We think this is the first report about the position of prickle with statistical data and mathematical model in the history of rose. Those data will help to understand the molecular mechanism of the development of prickle and to clear the role of prickles.

## Results

### Spatial pattern of prickles

*Rosa hybrida* cv. ‘Red Queen’ that has many prickles on the stem is used for the experiments (Fig. 1a). We measured the height and the angle of prickles during blooming the rose. We defined the measured height from the root of the shoot to leaf or prickle, *H*, and the measured angle of leaf or prickle with a stem as the axis, *θ* (Fig. 1b). The angle of leaves of *Rosa hybrida* cv. ‘Red Queen’ showed spiral pattern based on the golden angle Φ (Supplementary Fig. 1). *θ* was modified to *θ*_*c*_ that is cumulative value of *θ*, where *θ*_*c*_ = *θ* + 360*N* and *N* is non-negative integers, in order to express the relation of degree and height as a function (deep red line in Fig. 1c and Supplementary Fig. 2). From the low position leaf to the high position leaf, *N* of each leaf was selected so that *θ*_*c*_ of leaf increase monotonically. The spline curve connecting the leaf position *θ*_*c*_ is drawn on the *θ*_*c*_ − *H* plane (Fig. 1D (deep red line)). The spline curve is expressed as a function of height, 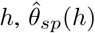. We defined the angle from the spline curve as *ϕ* (Fig. 1d).

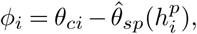

where *i* is the index of prickle, is assigned in order from bottom to top. *N* of prickle was selected so that *ϕ* of prickle ranged from −90° to 270°.

**Fig. 1.**
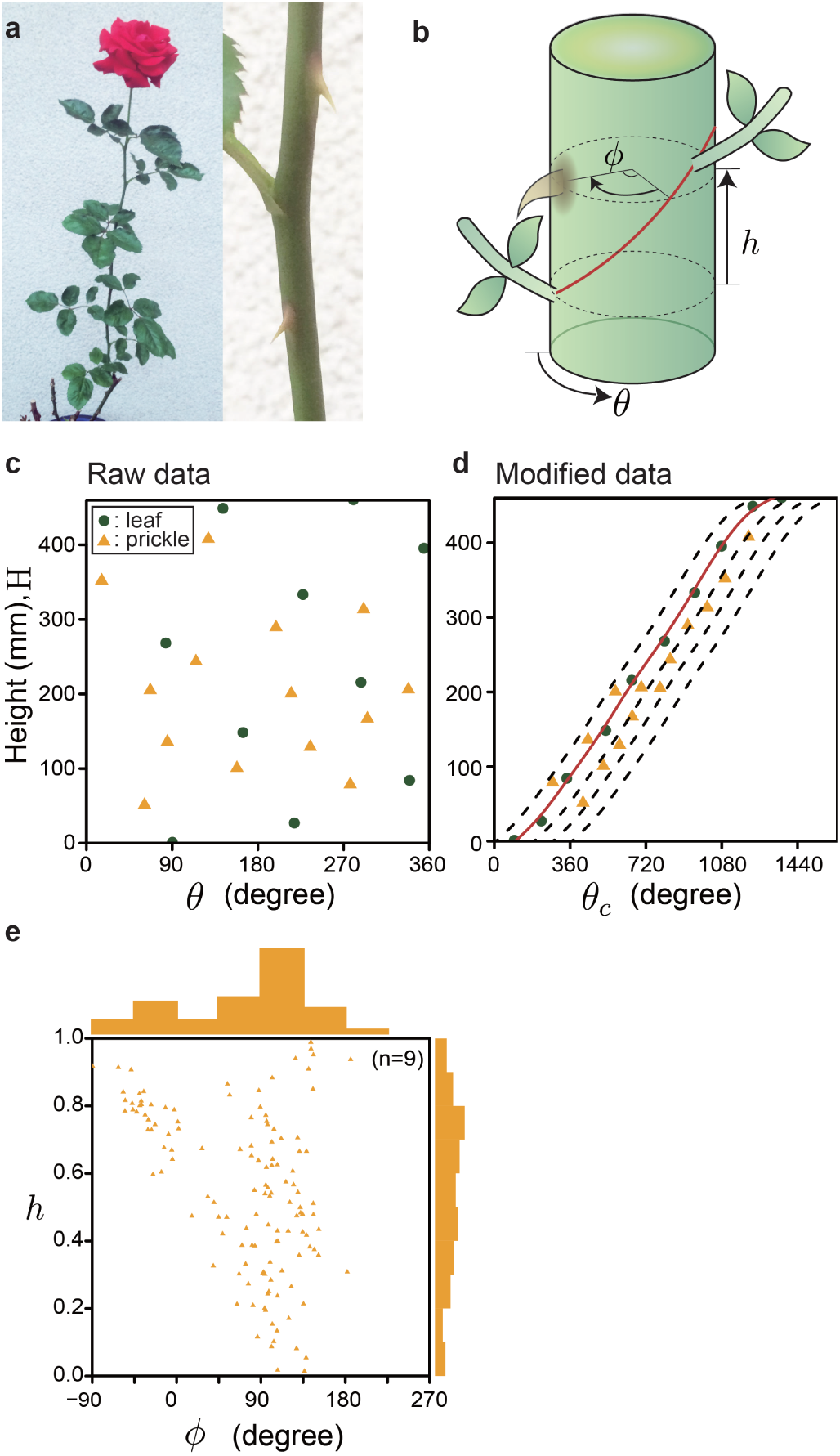
Pattern of prickles. a, Complete view and prickle of *Rosa hybrida* cv. ‘Red Queen’. b, *θ* is the measured angle of leaf or prickle. *ϕ* is the degree between prickle and the spline curve at same height on the stem. Let *h* define as the height of prickle between adjacent leaves, where the length between two leaves is 1. c, The position of leaves and prickles on the stem. Green circle is leaf. Yellow triangle is prickle. d, Deep red line show the spline curve connecting leaves. The dotted line was drawn by adding *−*90°, 90°, 180°, or 270° to the spline curve. e, Scatter plot of *ϕ* and *h* profile from measured samples (n=9). The histogram represents the distribution of *ϕ* or *h* from measured samples.

We discovered that the prickle occurs frequently in the range of *ϕ* between 90° and 135° at any *h* (Fig. 1E). If *ϕ* is between −90° and 0°, prickles were emerged at over *h* > 0.6. On the other hand, there are almost no prickles in other areas. Those results strongly demonstrate the position of prickle has a pattern.

### Mathematical model of the patterning

The fact that prickles emerge with a specific angle to leaves implies there are some regulations by primordia in the development of prickles. We constructed a simple model that reproduces the pattern of prickles on plane surface centered on apical meristem. The model assumed that periodically-generated primordia regulate the development of prickles through producing the inhibitor. Primordia moves straight away from the top of meristem. The direction of migration of primordia is shifted by Φ from the direction of the last produced primordia. The priming circle where the location of prickles is determined locates at a distance from the top of apical meristem. We also assumed the diffusion rate enough rapid that the inhibitor rapidly get equilibrium. Thus, distribution of inhibitor on the meristem is two-dimensional Gaussian distribution, implying that the concentration of inhibitor on the priming circle can be described as von Mises distribution, *k* exp(*m* cos *ϕ*), where *m* is a concentration parameter and *k* is a time-dependent parameter. These parameters, *m* and *k*, depend on time in our model. *m* should be proportional of the distance between the source of inhibitor and the center of the priming circle. Thus, we described the time dependence of *m* as *m*(*t*) = max [*αt* + *β*, 0], where *α* > 0, suggesting that primordia moves at a constant velocity. In addition, we hypothesized the production rate of inhibitor as a function of time, *k*(*t*) that maximize after when the primordia had passed through the priming circle. We assumed *k*(*t*) is a piecewise linear function that has three parameters *T*_*a*_, *T*_*b*_, and *T*_*c*_ (see Model in Methods). It imply that the position of prickles locating between *n*-th and *n* + 1-th primordia can be determined by the distribution of inhibitor produced by *n* + 1th, *n*th and *n* − 1th primordia (Fig. 2a).

**Fig. 2.**
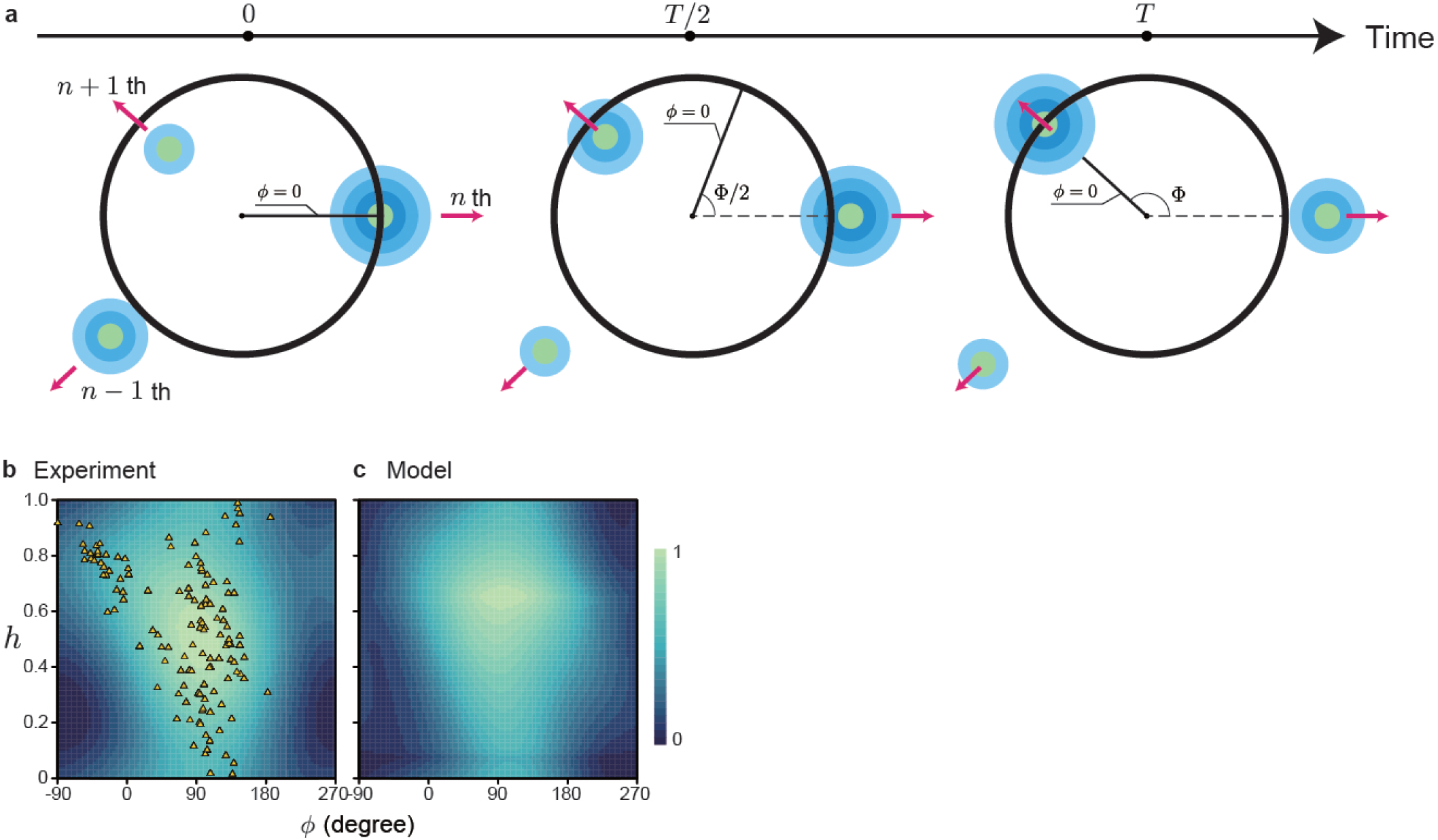
A model for development of prickles. a, Schematic diagram for distribution of inhibitor for development of prickles produced by primordia. Primordia are periodically created with a angle of Φ on a meristem. We assumed each primordia produces a inhibitor for creation of prickles and moves to outside at a constant velocity as growing. The prickles can be produced at a distance from the top of meristem (black circle) and the probability of synthesis of prickles on the circle is proportional to reciprocal of concentration of the in inhibitor. The position on the circle can be specified by *ϕ*, the angle from the spline curve. Black line from the center to the edge show the point of *ϕ* = 0 at each time. The amount of secretion of the inhibitor follows *k*(*T*). b, Estimated density of prickles on the *ϕ*-*h* plane through kernel density estimation for real data (Fig. 1e). c, Computer simulated density of prickles with a set of optimized parameters at *α* = 0.267, *β* = 0.139, *T*_*a*_ = *−*0.909, *T*_*b*_ = 0.980 and *T*_*c*_ = 1.590.

The parameter in the model, *α, β, T*_*a*_, *T*_*b*_, and *T*_*c*_ were optimized to maximize the correlation between the actual pattern of prickles, *p*(*ϕ, h*) and reciprocal of the amount of inhibitor, *f* (*ϕ, t*)^*−*1^. *p*(*ϕ, h*) was estimated from the real data (Fig. 2b and Methods). The optimized model qualitatively reproduced the actual distribution of prickles that peaks around 90°. Note that we repeated the optimization procedure with 10^4^ initial parameter sets. The parameters converged to either of two distinct parameter sets through the optimization. Fig. 2c was drawn based on the set of parameters that produces the distribution of *f* with the highest correlation to real data. The other set of parameters also produces the similar distribution of *f* although *k*(*t*) obtained from the parameters is shifted by approximately 0.2*T* (Supplementary Fig. 3). Additionally, We evaluate the dependency of the parameters relating to the diffusion of the inhibitor on maintaining the prickle patterning (Supplementary Fig. 4a). As a result, it was found that the distribution of the inhibitor being concave downward is crucial for reproducing the pattern of ‘Red Queen’ (Supplementary Fig. 4b, lower graphs). On the other hand, the pattern is relatively robust for quantitatively altering the distribution of the inhibitor itself (Supplementary Fig. 4b, upper graphs).

### Reproduce the another pattern on the other plant species

We tested that our model is able to reproduce the pattern of prickles in other plant species. Many plants show a pair of prickles under the root of leaf, not stipule (Fig. 3a). For example, *Rosa hirtula (Regel) Nakai* and *Acacia seyal* has a pair of the prickles at the same height (15) (Fig. 3a). We found the parameter set showing pair pattern on *ϕ*-*h* plane through the optimization algorithm (Fig. 3b, 3c). The releasing time of the inhibitor in *Acacia seyal* is different from ‘Red Queen’ (Fig. 3d). This result indicates differences in the parameter sets of the diffusion can cause the diversity of the prickle patterning in the kingdom Plantae (Supplementary Fig. 5).

**Fig. 3.**
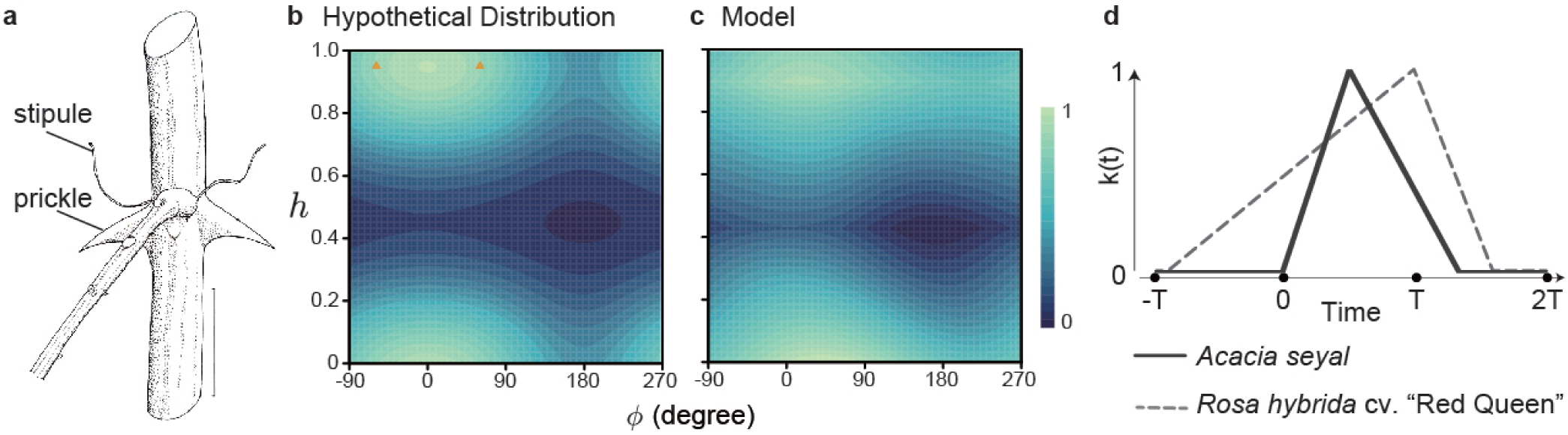
Estimation of the other pattern. a, Drawing of *Acacia seyal* which is modified from the Adrian’s book at page 146 and 147(15). b, The *h*-*ϕ* plane drawn by estimated prickle pattern of *Acacia seyal*. The *h* is 0.95. The *ϕ* of the two prickles is 60 and -60. c, Computer simulated density of prickles with a set of optimized parameters at *α* = 0.227, *β* = *−*0.144, *T*_*a*_ = *−*0.023, *T*_*b*_ = 0.489 and *T*_*c*_ = 1.330. d, Plot of the time dependent production rate *k*(*t*).

### Interaction between prickles

Here, We have focused on the interactions between primordia and prickles so far, but interaction between prickles was indicated from the length *d*_*pp*_ between a prickle and the nearest one. The *d*_*pp*_ of the ‘Red Queen’ showed the growing prickle strongly inhibit the development of other prickles within about 5 mm radius (Supplementary Fig. 6a). It indicates growing prickles prevent the growth of other prickles surrounds growing one, like excluded volume effects. If there is no excluded volume effects and the position is determined randomly, fused prickle must be more appeared on the stem (Supplementary Fig. 6b).

## Discussion

In future studies, the prickle position on the other species of roses should be measured, in order to confirm that the patterning based on our model is conserved between the species. The number of prickles on the stem has a large variation between species of roses. Those differences of prickle patterning could be reproduced by the changing parameters. Although analysis of entire genomes of roses is few compared with those in other species and because sequencing remains challenging(16, 17), comprehensive analysis of genome and the pattern will help to detect genes that cause the differences. In addition, time-dependent development of prickles should be observed in future studies. Small prickles whose height is lower than 1 cm were observed (Supplementary Fig. 7). Especially, they appeared frequently at the bottom side of the stem. A similar phenomenon has been reported on the stem of Rosa hybrida cv. ‘Laura’(18), but the patterning of the small prickles is not understood. It is also not known whether the small prickle grows into the mature prickle. The observation may also reveal the Φ distribution of the prickles is slightly changed depends on the height (Supplementary Fig. 8a). We found, when dividing the height from the root to the top leaf into the three layers, the unimodal distribution is shown at the top layer, but bimodal distribution is shown at the other layers (Supplementary Fig. 8b). Those facts may indicate the parameters of the prickle slightly changes as the rose grows.

It is generally accepted that spiny structures on the stem play a key role in defense against herbivore(19, 20). In addition, prickle is considered as attachment devices to prevent slipping off the axes from supporting structures. Previous studies have discussed the two roles by focusing on the shape, physicochemical and biological properties of the spiny structures on the stem(21–23), but herein we shed light on the possibility that the spatial pattern of the prickles makes the two roles more effective. Spiral patterns of leaves are based on the golden angle which avoids overlapping of growing leaves when viewed from the top of the stem, thus increasing the efficiency of photosynthesis. Prickles occurred frequently at the specific angular position relative to the spiral curves of leaf position. Therefore, the prickles also appear in the spiral pattern which avoids the overlapping like a leaf (Supplementary Fig. 2). The character of the pattern would perform more efficient at preventing slipping off and at defense from an herbivore.

Spiny structures on the stem of plants are derived from many types of tissue of plants. Although, the prickle on the stem of rose is derived from epidermal cells on the stem, it is known that the spiny structures on the other plants, such as *Robinia pseudoacacia* and *Zanthoxylum piperitum*, are derived from stipule. Interestingly, the position of prickles of *Z. piperitum* shows the pair pattern like Figure 3a, but *Z. schinifolium* loses the pair pattern(24). The pattern looks random and is not determined yet. It is not revealed that why the difference of spatial pattern of spiny structure derived from stipule happens. In the future study, our model of prickles may be used to explain the difference of spatial pattern of spiny structures derived from stipule.

Although humans have characterized roses in numerous reports since the start of cultivation several thousand years ago, this is the first report to show the spatial pattern of prickles with the statistical data and to suggest the mathematical model. Moreover, in this paper, we suggest the spatial pattern will affect the role of prickles. Further investigation will clarify the role of the patterning and reveal the entity of the diffusing factors. Even though herein we suggest the simple model, further developing of our model will help to build a novel theory of plant development based on diffusion, especially the protrusion structures on the epidermis of the plant.

## Methods

### Plant materials and measurement

*Rosa hybrida* cv. ‘Red Queen’ was purchased from Keisei Rose Nurseries (Chiba, Japan). The plants were cultivated in pots placed at the open field under natural daylight. The angle and position of prickles and leaves on the lateral axis were measured by protractor and calipers. The direction from the lateral axis to the main axis was set to 0°. Small prickles with a height of 1 cm or less were not measured. Clockwise and counter clockwise spiral pattern of leaf were not distinguished in this study. All data was analyzed by R program ver. 3.5.2.

### Model

We assumed that the emergence of prickles can be inhibited by the diffusive molecules secreted from the primordia that radially move down from meristem. The primordia can be produced with a constant divergence angle Φ at every time interval *T*. The priming zone where the prickles are emerged locates on the meristem. If the priming zone is not large, the zone can be simply represented as a circle. Let *ϕ* be an angle on the circle of priming zone and set *ϕ* = 0 for the direction of *n*th primodia, without loss of generality. Suppose that the inhibitor can diffuse on only the surface on the meristem. Concentration of diffusive particles produced in a single point and randomly diffuse on a 2D plane obey the normal distribution at equilibrium state. Then, the particles on a circle with a distance *m* from the source are distributed as a von Mises distribution (25). Thus, the intensity of inhibitor at a angle of *ϕ* on priming circle can be approximately proportional to von Mises distribution, i.e., *f*_*i*_(*ϕ, t*) = *k*(*t*) exp(*m* cos(*ϕ* − *ϕ*_*i*_)), where *ϕ*_*i*_ is the direction of *i*th primodium and *k* is a time-dependent parameter. We here set *t* = 0 at the time when *n*th primoidia pass the priming zone and *m*(*t*) = max[*αt* + *β*, 0]. In addition, we assumed the secretion of inhibitor arises when the primodia is passing through vicinity of priming zone, thus depends on The magnitude of secretion, *k*(*t*), was represented as a piecewise linear function:

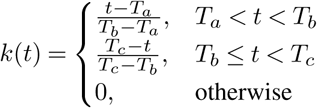

where *T*_*a*_ < *T*_*b*_ < *T*_*c*_, *T T*_*a*_, 0 < *T*_*c*_ ≤ 2*T*, implying only 1, *n* and *n* + 1th primodia contribute to distribute the inhibitor on the priming circle. Taken together, the total intensity of inhibitor produced from every primodium at time *t, f*(*ϕ*) can be described as,

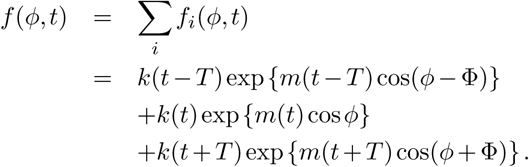

The parameters in this model, *α, β, T*_*a*_, *T*_*b*_ and *T*_*c*_ were chosen to reproduce the experimental data through optimization algorithm. The real distribution based on the observation, *f*_*r*_(*ϕ, t*) was obtained through kernel density estimation based on the observed arrangement of prickles. We sampled the values of *f* (*ϕ*_*i*_, *t*_*j*_) and *f*_*r*_(*ϕ*_*i*_, *t*_*j*_) where *ϕ*_*i*_ = 2*πi/N*_*d*_, *t*_*j*_ = *jT/N*_*d*_ and division number *N*_*d*_ is 100. The cost function was introduced as the Pearson correlation coefficient for the sampled values of *f* and *f*_*r*_, multiplied by *−*1. Kernel density estimation was performed by the function kde2d in the R MASS Package ver 7.3-50. Optimization procedure was performed by the function optim with BFGS method in the R stats package ver 3.6.0.

## Acknowledgements

We thank to Prof. Hirokazu Tsukaya (Univ. of Tokyo) and Assoc Prof. Munetaka Sugiyama (Univ. of Tokyo) for valuable comments to our research, R. Yasui (Kyushu Univ.) for helping drawing an illustration of rose of Fig. 1b and Sergey Melnikov (Yale Univ.) for many valuable comments to our manuscripts. Fukuoka City Botanical Garden where the idea of this paper first came up for us. Rose garden in Jindai Botanical Garden and Chiba Kashiwanoha Park where the idea was sophisticated.

## Author contributions statement

Cultivation study and measurement were performed by K.A. Analyzing data and writing manuscript was performed by K.A. and H.I.

## Supplemental Information

**Supplementary Figure 1.**
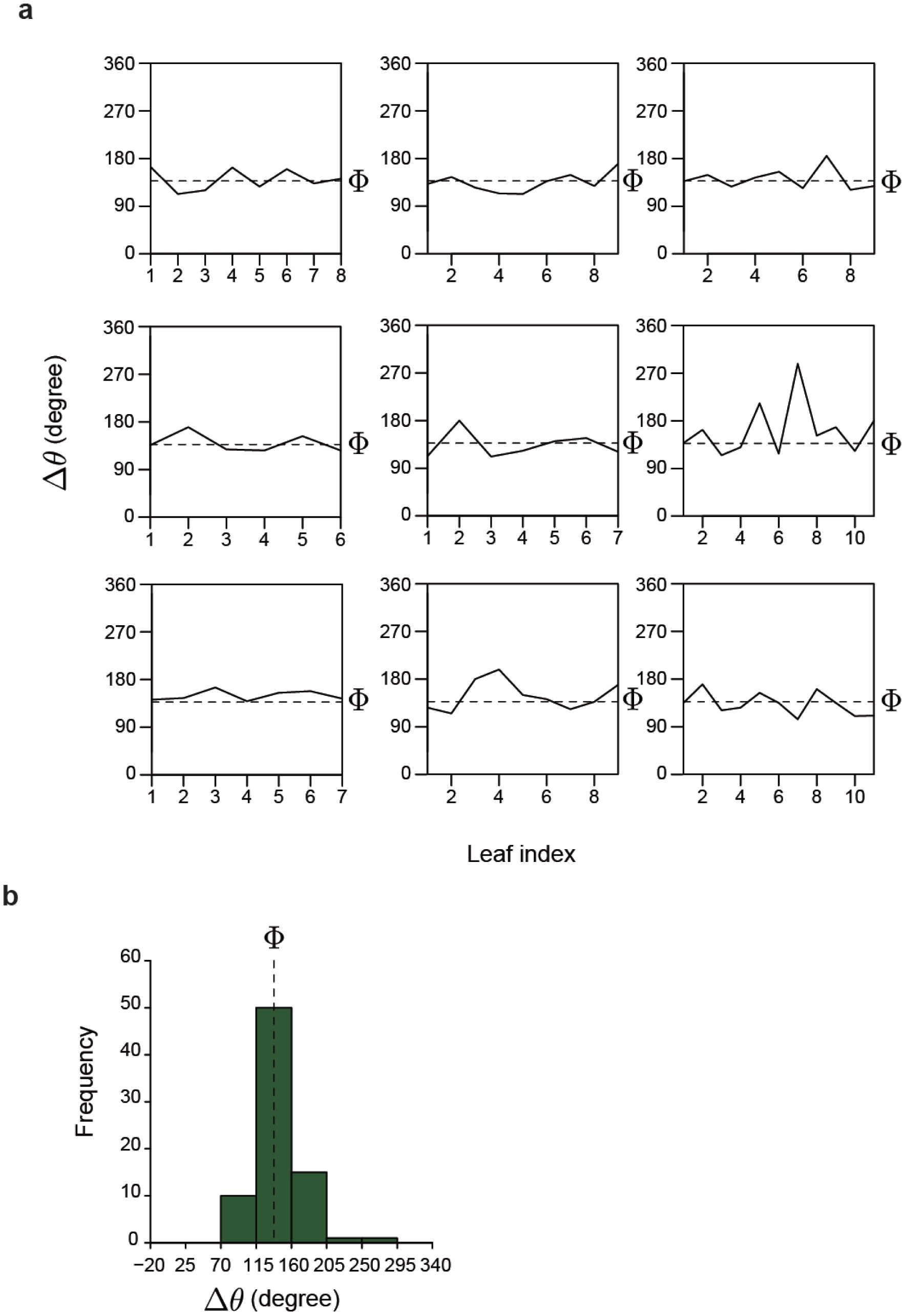
Differential values of leaf degrees. a, In order from the top of each stem, the number of leaf index was assigned. Δ*θ* is the difference of the *θ*_*c*_ between the adjacent leaves. The Φ is about 137.5° which in golden angle. b, The histogram represents the Δ*θ* from all samples.

**Supplementary Figure 2.**
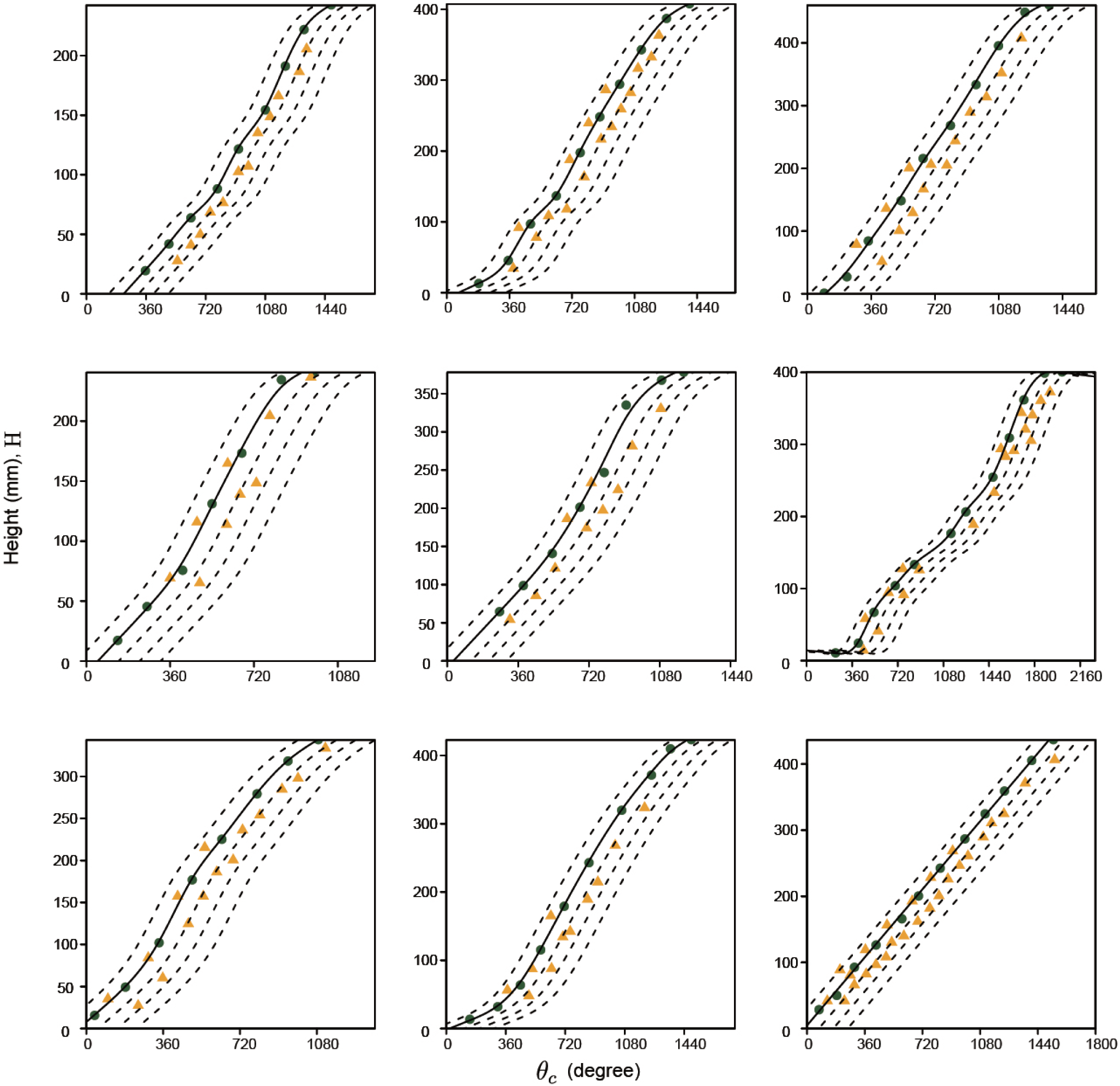
The *H*-*θ*_*c*_ plane of all samples.

**Supplementary Figure 3.**
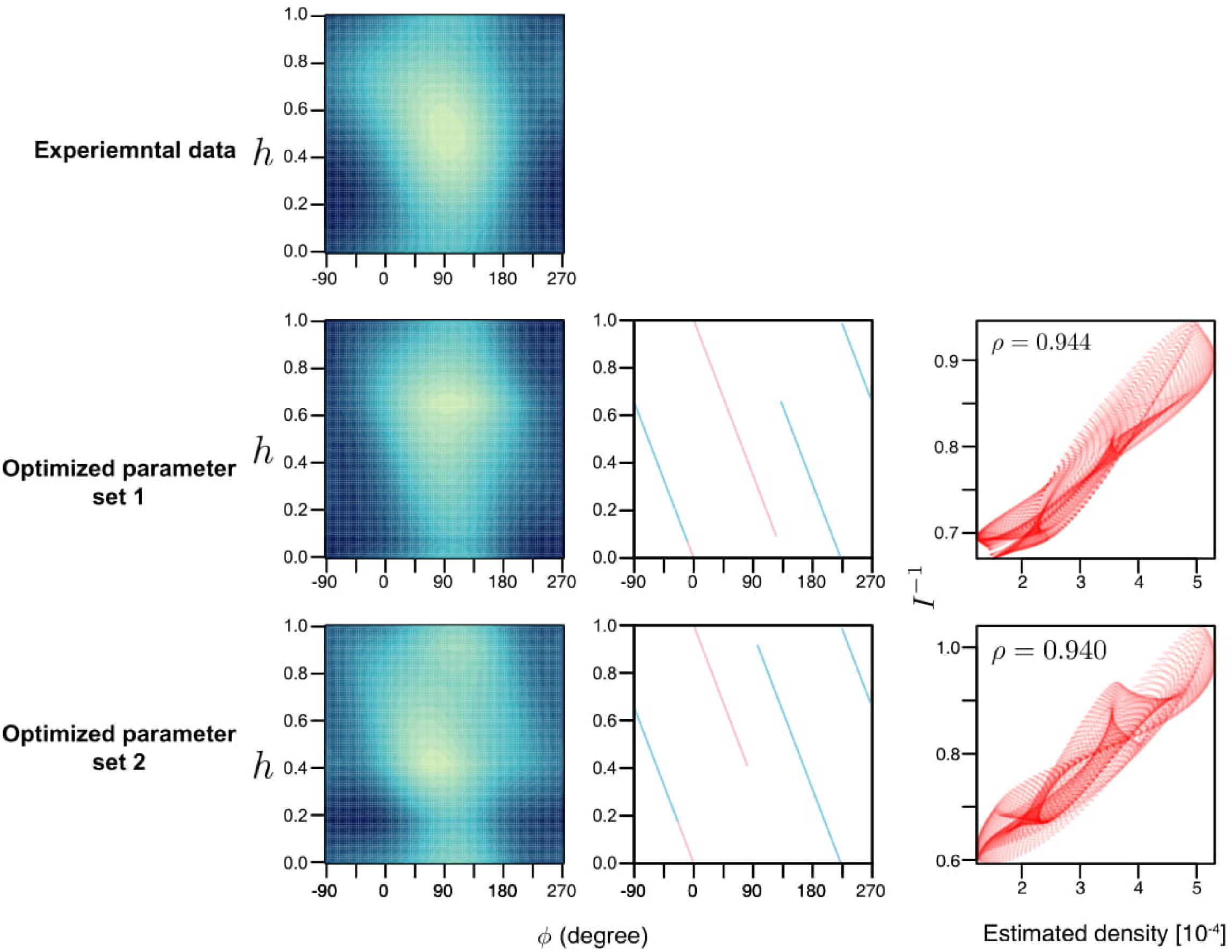
Analysis variation of *H*-*ϕ* plane using optimized parameter. a Estimated distribution of prickles on the *ϕ* -*h* plane through kernel density estimation for real data (the same figure as Fig 2b). Two parameter sets were obtained by maximizing the Pearson’s correlation between density from real data and *I*^−1^ calculated from a model. b, e distribution of *I*^−1^ obtained from the optimized model with each optimized parameter sets. c, f The trajectory of the primordia are drawn on the *ϕ* -*h* plane. The red and blue represents the interval where the amount of secretion of inhibitor are increasing 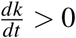 and decreasing 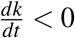, respectively. d, g The correlation between the estimated density obtained from real data and *I*^−1^ obtained from the optimized model. The correlation coefficient *ρ* is displayed in each figure.

**Supplementary Figure 4.**
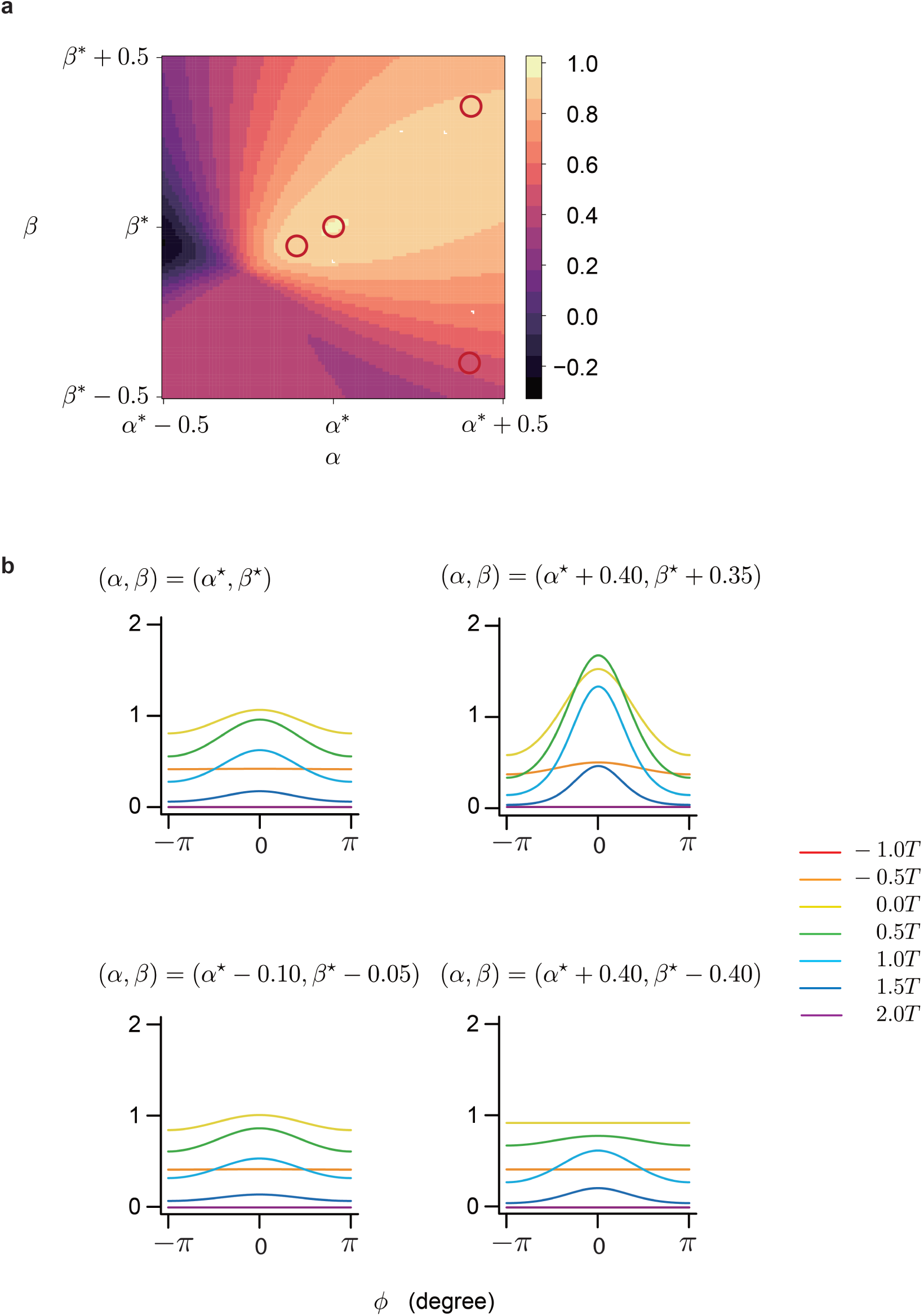
Parameter sensitivity. a, The correlation between the real data and the simulation data. The *T*_*a*_, *T*_*b*_ and *T*_*c*_ is fixed at optimized parameters at Figure 2c. *α*^∗^ and *β*^∗^ is 0.267 and 0.139. b, The *f* (*ϕ, t*)-*ϕ* graph is drawn at each point.

**Supplementary Figure 5.**
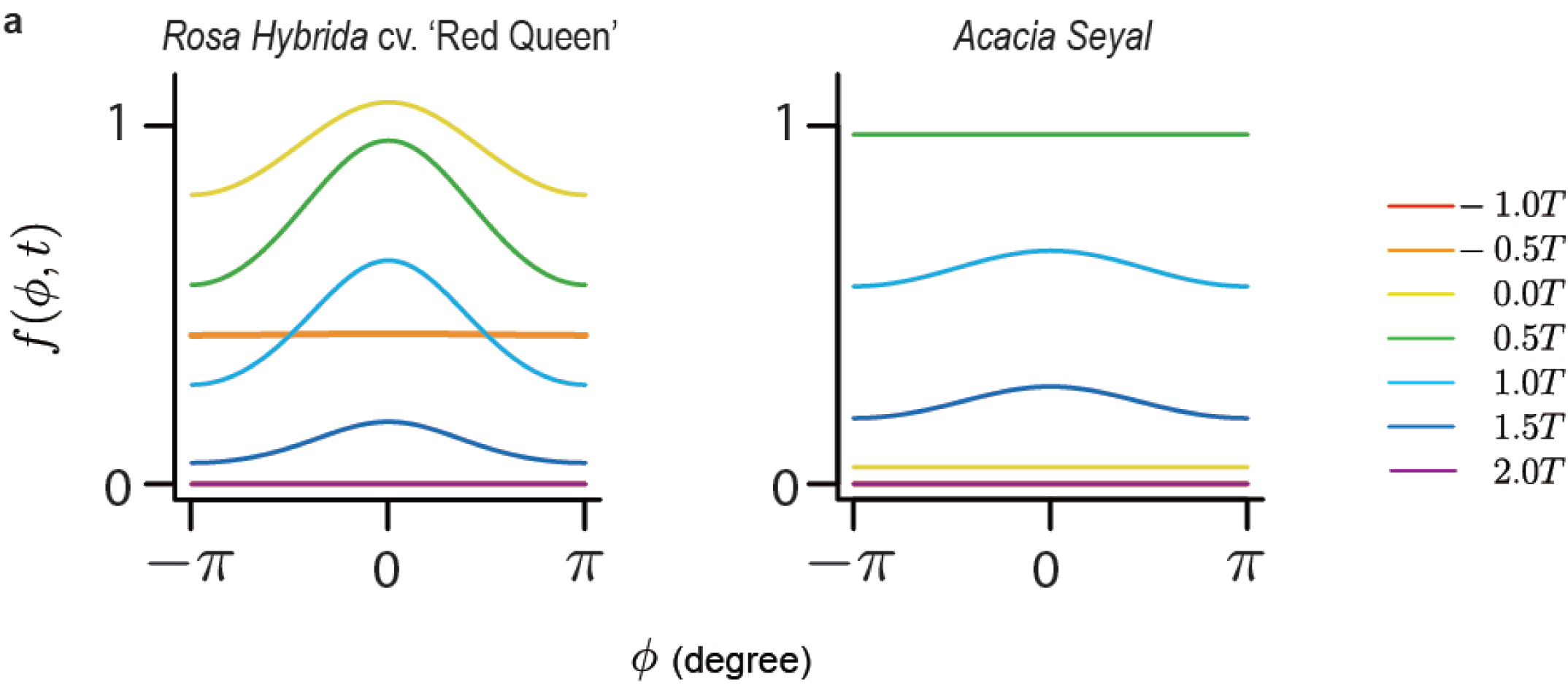
*f* (*ϕ, t*)-*ϕ* graph. a, The graph drawn by the set of parameters of Figure 2c. b, The graph drawn by the set of parameters of Figure 3c.

**Supplementary Figure 6.**
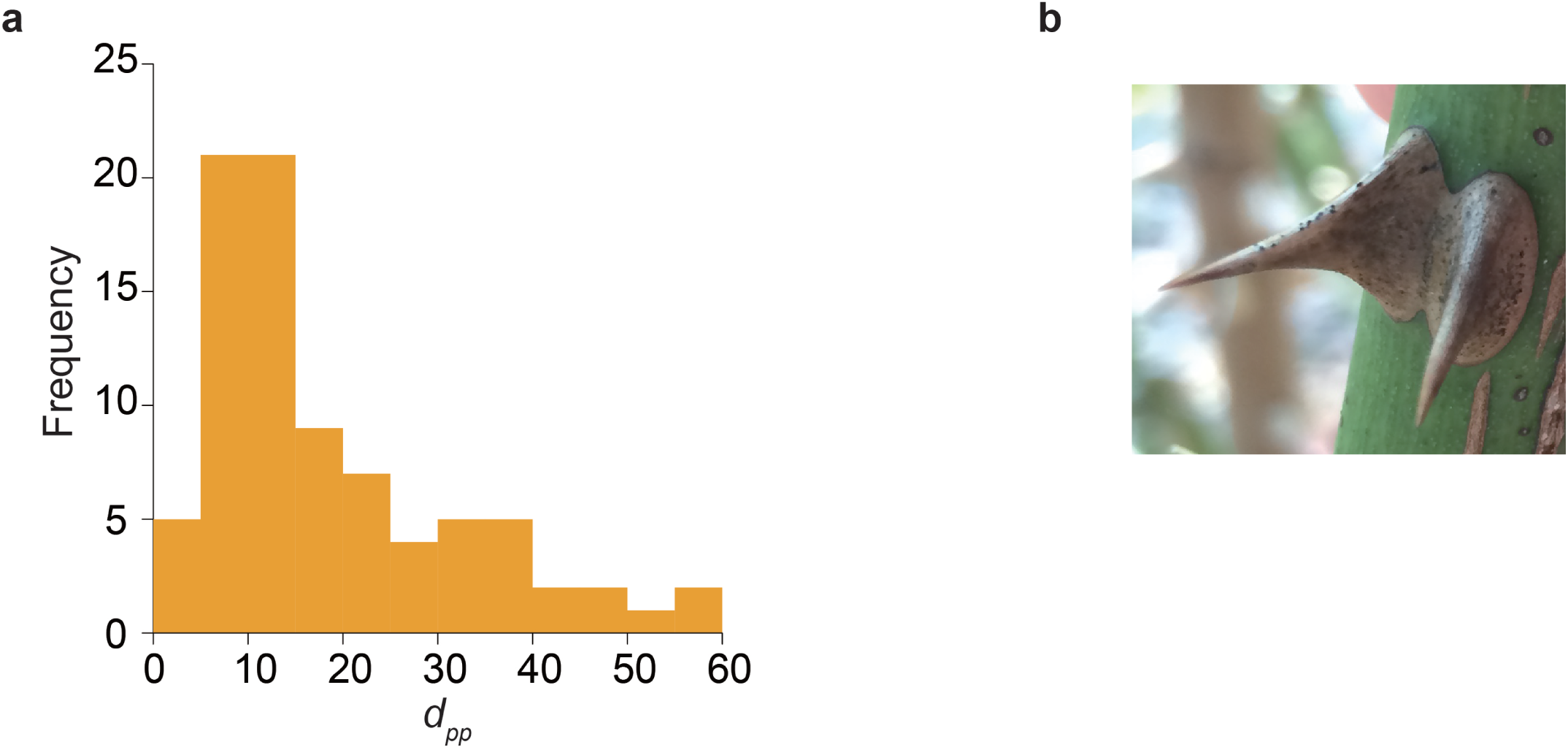
Interactions between prickles. a, the distribution of *d*_*pp*_. b, Fused prickle on the stem of *Rosa hybrida* cv. ‘Carinella’.

**Supplementary Figure 7.**
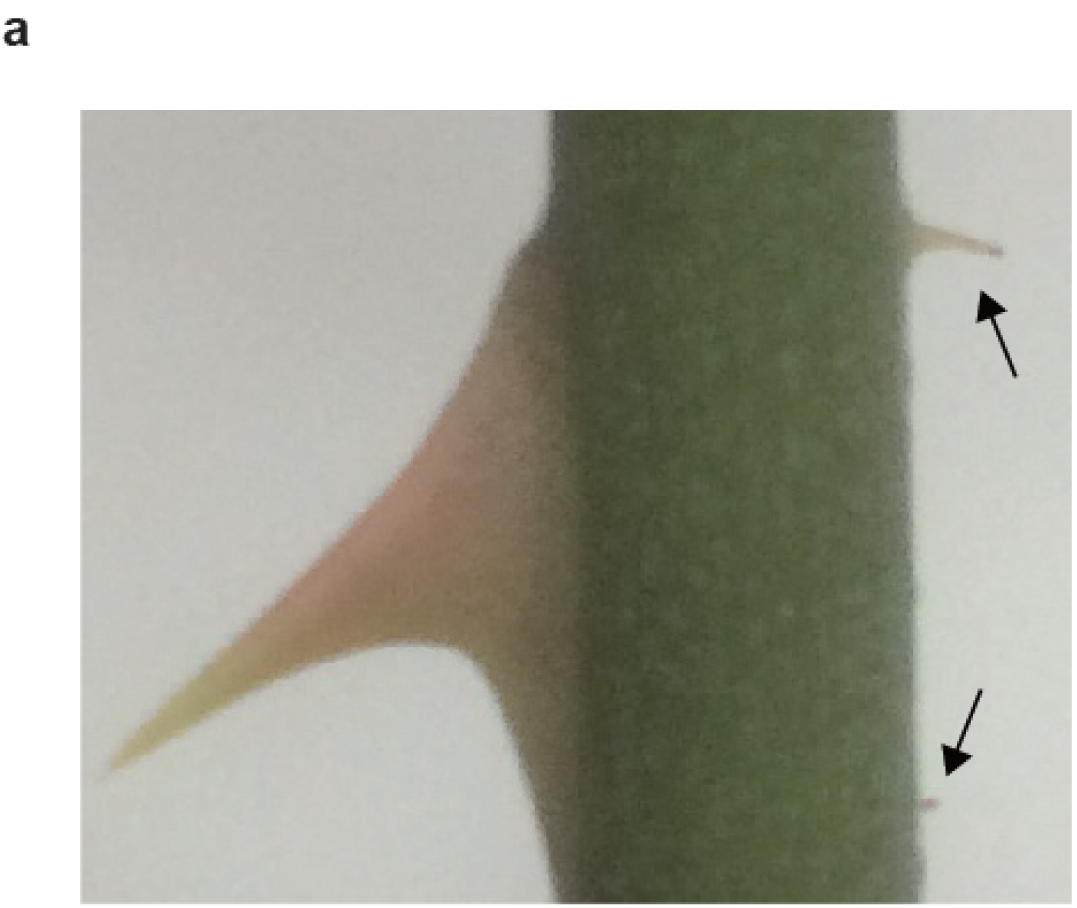
Small prickles. a, Large and small prickle on the stem of *Rosa hybrida* cv. ‘Red Queen’. The black arrow is pointing to the small prickle.

**Supplementary Figure 8.**
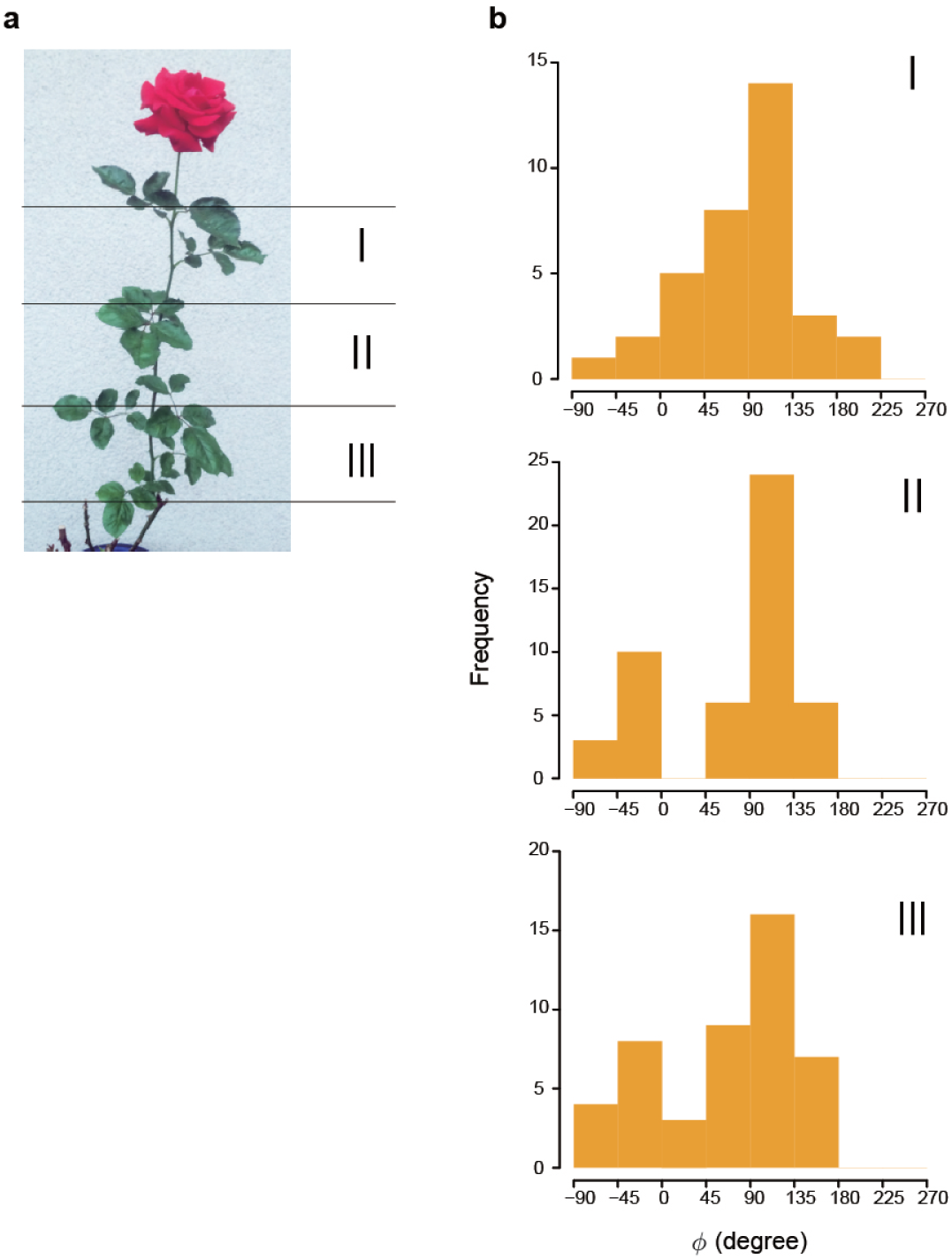
Bimodality of the distribution of *ϕ*. a, The divided area in height dependence. b, The histogram of *h* in the each area.

